# Effect of ethanolic extract of Cyperus rotundus reduces the surgical induced secondary lymphedema and oxidative stress in adult mice tail

**DOI:** 10.1101/2023.09.18.558373

**Authors:** Nikhil Pandey, Priyanka Mishra

## Abstract

**Background:** Lymphedema is clinically manifested as swelling in the extremities due to abnormal accumulation of protein rich in the extravascular interstitial space resulting in irreversible structural changes to the limb (s). The aim of this explorative work was to evaluate the effect of *Cyperus rotundus* root (CRR) ethanolic extract in a mouse tail model of secondary lymphedema. Method: Mice were temporally anaesthetised under sterile condition and the skin from the tail was removed from distal part of the trunk after leaving 1cm of distance. The animals were divided into Experimental control (EC) and *Cyperus rotundus* root (CRR) 80 mg/kg b.w. /day) treated groups. Change in tail volume and circumference were monitored for 20 days. TNFα, SOD and catalase were estimated from blood obtained through retro-orbital at day 0, 5, 10 and 15. Further TS of upper part of the tail skin was stained with H&E stain to document histological changes mRNA level of COX-2 was estimated from the skin.

**RESULTS:** EC group showed gradual rise in tail oedema post-surgery (PS), elevated inflammation, oxidative stress and histopathological alterations. However in CRR (80 mg/kg b.w./day) treated group shown the reduced tail oedema after post-surgery. TNFα, SOD and catalase levels were significantly less in CRR then EC group supporting anti-inflammatory potential. CRR protected the tail from structural damage. It also down regulated the expression of COX-2 in comparison to EC group. Conclusions: CRR ethanolic extract significantly attenuated secondary lymphedema indicating it potential for therapeutic use.

**Graphical abstract:** 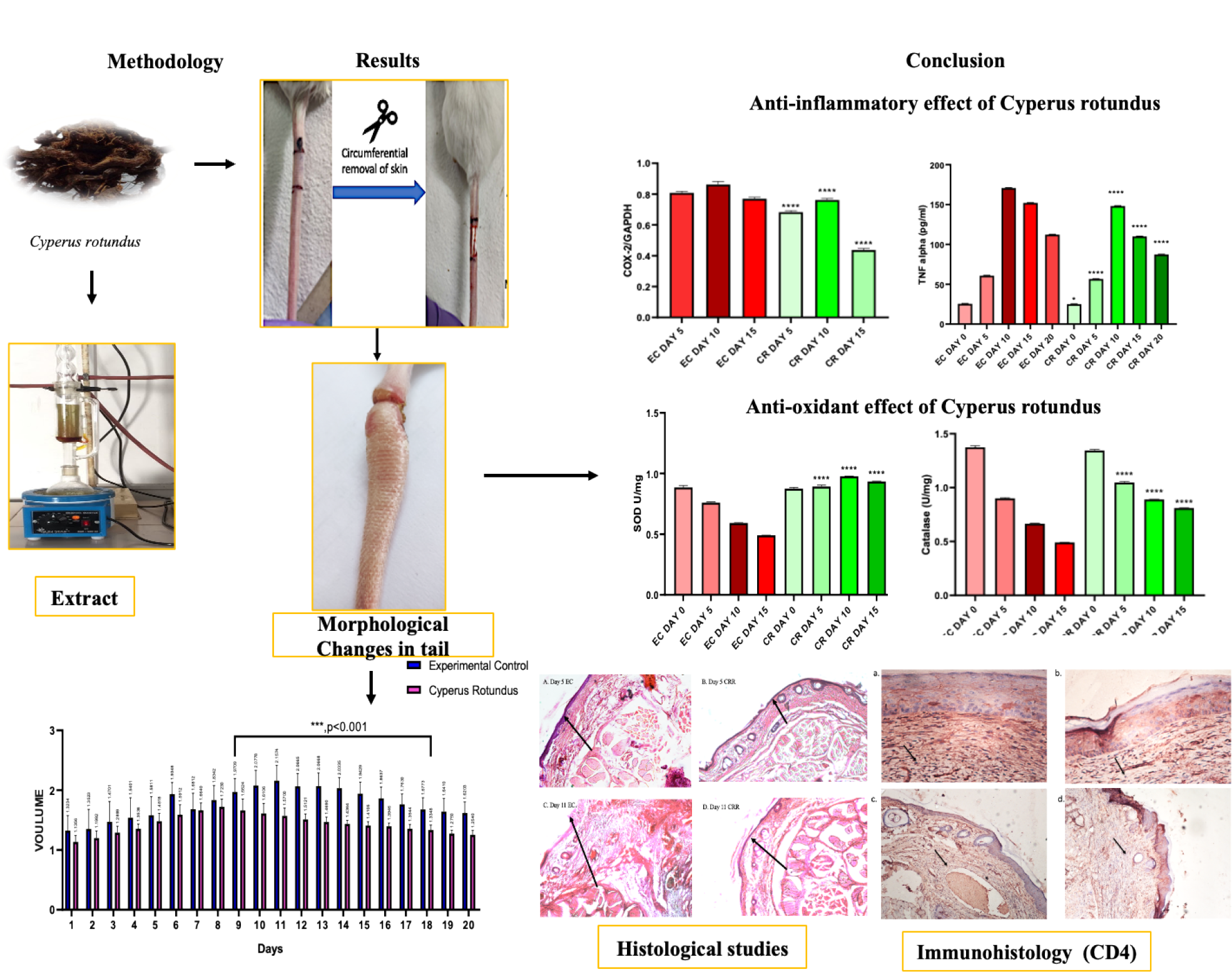

## 1. Introduction

More than 130 million people are affected with secondary lymphedema (SL), a chronic condition characterised by the accumulation of fluid, proteins, and adipocytes in the interstitium, resulting in the swelling of the affected limb.(1)(2) .The vascular network in the lymphatic system is vital to maintain the homeostasis of fluid in the tissue in order to maintain the tissue fluid homeostasis and to conciliate the local inflammation and the immune response. As the anatomical and the functional integrity of the vasculature in the lymphatic system is impaired, leads to the loss of fluid transport potential resulting to the condition of lymphedema which is the easily recognizable characteristics (3) .It is characterised by progressive swelling due to disrupt local lymph transport, leading to the permanent and disoriented changes in the tissue architecture. Due to surgical intervention or by infection, people develop the secondary lymphedema which gets worse in gradual manner mostly in developed countries. As SL is common and leads to loss of limb function due to vascular insufficiency, still lacks pharmacological treatment. (4).Hence at present the use of the pharmacotherapy agents in clinical and experimental studies are focusing on inhibiting inflammation. Earlier, approaches were made to reduce the chronic lymphedema through inhibiting the NfkB for example sodium selenite shown to reduce the lymphedema volume by directly inhibiting the expression of adhesion molecules and NfkB, But its further trial are needed to confirm these results. (6). Therefore the finding new therapeutic option is absolutely necessary. In the present work, we chose *Cyperus rotundus* (CR) which is known for its TNFα inhibitory activity (7) and is employed for its preventive action of the reduction of obesity and the reducing of the blockage in vascular system. (8).CR has been earlier reported to inhibit inflammatory cascade.(9) Therefore, we hypothesised that CR could be used in the management of SL by assessing its activity on inflammatory cytokines, oxidative stress and on histological changes observed in mice. Here based on the earlier work of Olszewski’s lymphedema model we chose the moue tail model to induce SL (10)(11).

## 2. Material and Methods

### 2.1. Material: Characterization of extract : Anti-bacterial analysis : Cytotoxicity Study

#### For surgical procedure

Carbon steel surgical blade No.23, Stainless steel split ring tweezer Ethanol (70%), sterile cotton, digital vernacular calliper(Mavotank Pvt ltd),For biochemical procedure-Mouse TNF alpha ELISA kit was purchased from Elabscience (E-EL-M3063), TRIzol (Ambion),TURBO DNA-free Kit (Invitrogen),RevertAid First Strand cDNA Synthesis kit (Thermo scientific).

### 2.2. Preparation and standardization of the extract

*Cyperus rotundus* roots (CRR) were procured from the botanical garden and its authenticity was verified by Centre of Advanced study in Botany, Institute of Science, BHU (voucher no. Cypera.2021/4). The 200g roots were powdered and then exhaustively extracted with ethanol in continuous Soxhlet extractor for 36 hrs. The ethanol filtrate was evaporated to a dryness using a Buchi RE11 rotavapor and Buchi461 water bath. The yield was 17.5 percent i.e 17.5 gm of crude Cyperus rotundus roots extract (weight of dried extract/weight of the powdered root). Fresh solution of the crude CRR extract was prepared by dissolving a given quantity of the ethanol extract in a small volume of tween 20 and made up to appropriate volume with the sterile water The Ethanol CRR extract was given in the dose of 80 mg.kg/BW/day orally to the mice of the CRR group up to twenty days. The experimental control group was given 1ml of drug vehicle (i.e 20% of Tween 80 in the appropriate volume with the sterile water).

### 2.3. Characterization of extract

### 2.4. Anti-bacterial analysis

The bacterial isolates Escherichia coli and Staphylococcus aureus were isolated from the lab. Bacterial cultures were maintained on nutrient agar (NA) plates and slants. They were sub cultured every 2 weeks and subsequently stored at 4°C. Inoculum Preparation Nutrient agar broth was inoculated with freshly sub cultured bacteria and incubated at 37°C for few hours. Such prepared inoculums were used to spread onto agar plates using sterile cotton to make a lawn of bacteria. Preparation of samples in different chemical.10 grams of sample was mixed with 40 ml of different chemical such as chloroform, water and ethanol. Sample mixed chemical was boiled in a boiling water bath for 30 minutes, cooled at room temperature and filtered through whatsmann No .1 filter paper and collected in a sterile container for further use. Extracts was kept at 4°C to preserve the antibacterial property before they were used for well diffusion assay with antibiotic disc. Antibacterial assay A loopful of each bacterial isolates were prepared in agar plates using a sterile cotton swab from the inoculums. The plates were dried for 15 minutes in a laminar air flow chamber. Then make a small well on each agar plates. The antibiotic disc was commercially available Antibiotic disc were used as the control and placed on the centre of the agar plates. Added 20 microlitre of extracts of samples on each well. All plates were incubating at 37°C for 18-24 hours and the zones of inhibition (diameter in mm) were measured on the agar plate.

### 2.5. Cytotoxicity Study: Cell lines

#### a) MCF7

Human Breast adenocarcinoma cell line (From NCCS, Pune), Cell culture medium: DMEM-High Glucose -(#AL111, Himedia), Adjustable multichannel pipettes and a pipettor (Benchtop, USA), Fetal Bovine Serum (#RM10432, Himedia), MTT Reagent (5 mg/ml) (# 4060 Himedia), DMSO (#PHR1309, Sigma), Camptothecin (#C9911, Sigma), D-PBS (#TL1006, Himedia), 96-well plate for culturing the cells (From Corning,USA), T25 flask (# 12556009, Biolite -Thermo), 50 ml centrifuge tubes (# 546043 TORSON), 1.5 ml centrifuge tubes (TORSON), 10 ml serological pipettes (TORSON), Centrifuge (Remi: R-8oC). Inverted Biological Light microscope (Biolinkz), 37°C incubator with humidified atmosphere of 5% CO2 (Healforce, China)

### 2.3. Animal and experimental design

The experimental protocols were approved by the Animal Ethical Committee of IMS-BHU (letter No. Dean/2021/IAEC/2567). Inbred male mice Swiss albino strain (15-20gm) was purchased from the Central Animal facility. They were acclimatized for 7 days in ambient conditions of temperature, and a day-light cycle of 12 hrs. normal room temperature with 45%-55% of the humidity. The normal diet was provided during the course of the whole experiment. Animals were divided in the following group as shown in (Table 1)

**Table 1.** -Experimental design of the study.

| S.No. | Group | Intervention | Route |
| --- | --- | --- | --- |
| 1 | I- Experimental control (n=30) | 20% of Tween 80/day | Per oral |
| 2 | II- Treatment (n=30) | CRR (80 mg/kg b.w/day) | Per oral |

### 2.4. Model to induce SL

We carried the study for secondary lymphedema in the tails of Swiss albino mice (20-25gm). The tail skin was removed after leaving 1cm of distance from the base of the trunk. Cut was introduced in sterile condition and no vessels were damaged. The administration of ether was applied for the temporary anesthesia and skin and subcutaneous tissue between from the base of 5-10 mm to the distal region of the tail was removed (fig 1). The mice were kept in separate cages to avoid the damage being caused by the other mice. We measured the volume of the tails every day for 20 days, thickness and histological studies were also done to assess the lymphatic adequacy. The tail was ethically amputated in aseptic condition on due time period for the examination of tail for mRNA expression and histological studies for its gross examination of the tail. During the given time scale the photography of the tail was carried out to examine the change in inflammation in lower and upper part of the cut. The measurement of tail was done with digital caliper.

**Figure 1:**
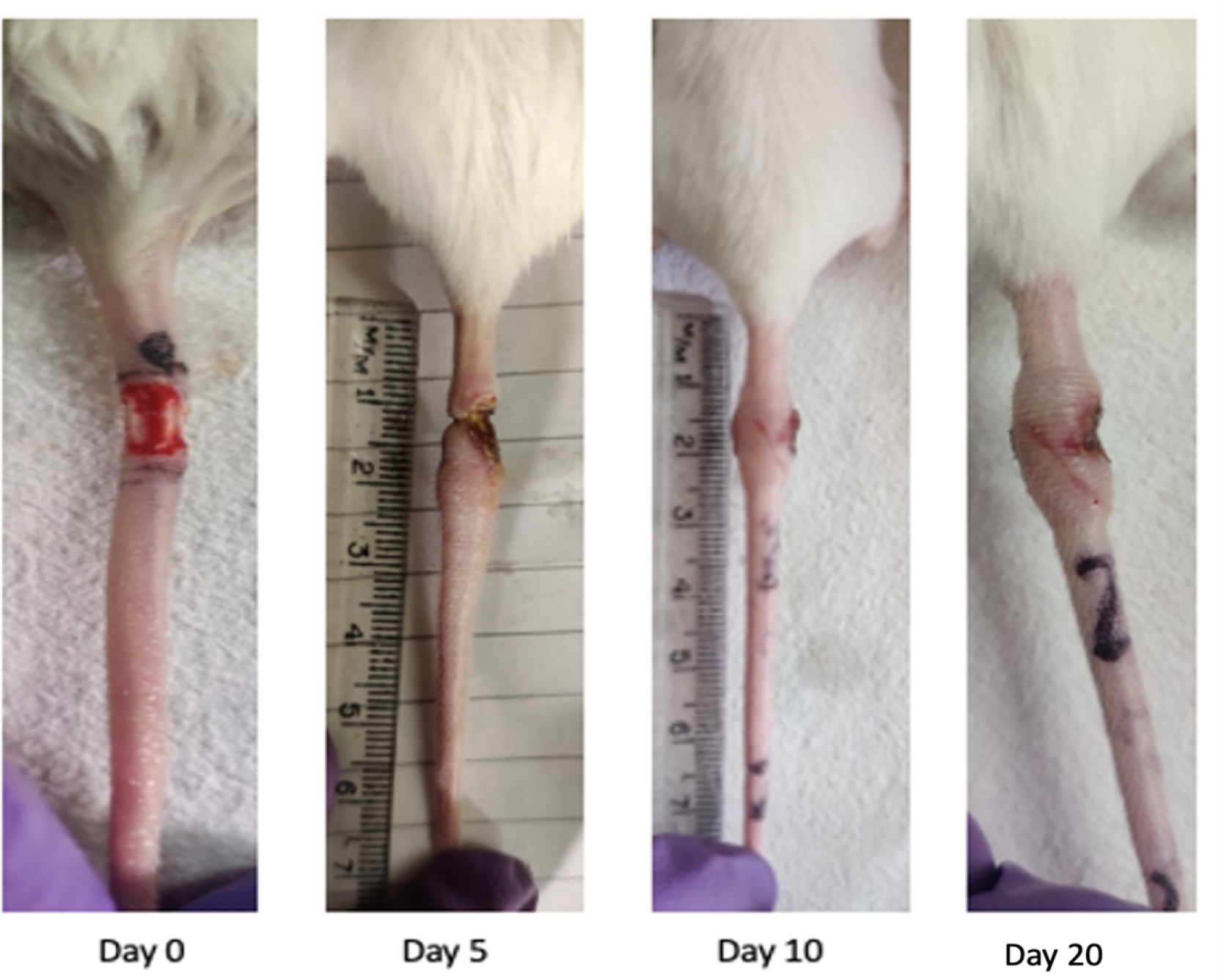
Morphological assessment of mice tail at day 0,5,10 and 20 in EC group.

### 2.5. Parameters studied

#### 2.5. A. Measurement of morphological changes and tail oedema after post surgery

Mice were analyzed visually for assessing morphological condition on day 0, 5, 10 and 20 for both groups. Tail Odema were measured in terms of volume and circumference measurement followed from previous study (12).

##### Determination of Volume measurement at inflamed area

At post -surgery every mice were taken for the measurement of tail diameter of by digital vernacular calliper of upper and lower ends of the swelling. Along with the a ruler for measuring the length, digital vernacular calliper was used to find the thickness increase and decrease in the murine tail. Photos were taken to measure the diameter of each region .Besides after converting the diameter into circumference by using mathematical equation :

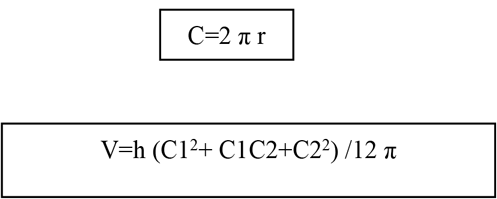

Finally the volume was measured by using the following mathematical equation where tail was assumed to be a sliced cone. (13).

#### 2.5. B. Measurement of biochemical changes

##### Determination of TNF-α level

Wounded part of the mice tail were surgically removed at day 0,5,10 and 15 was homogenised in cold PBS, centrifuged and supernatant was kept for TNF-α estimation through a procedure stated by Elabscience mice ELISA kit (E-EL-M3063).

##### Determination of oxidative stress

Part of the inflamed area from the upper part of the wound was taken day 0,5,10 and 15 and placed in 1ml of PBS and used for further estimation. Superoxide dismutase (SOD) activity was quantified by Beauchamp method where the rate of inhibition of nitro blue tetrazolium (NBT) reduction in the presence of riboflavin as described (14).The catalase enzyme activity was measured by the Aebi’s method by observing the H_2_O_2_ breakdown at the 240nm (15).

#### 2.5. C. Measurement of histological changes

The collected tail samples at day 5 and 11 were removed surgically from both groups. It was fixed in formaldehyde and was dehydrated, embedded in paraffin, sectioned into slices (5 μm) and stained with haematoxylin and eosin (H&E).The Samples were observed at 10X magnification.

##### D. Measurement of mRNA expression of COX-2

Inflamed upper part of the tail was taken for the total RNA extraction on day 5, 10 and 15 from both the EC group and CR group. As per the user’s manual, dissolved RNA (DEPC-water) was quantified through spectrophotometer. 1mg of tissue was subjected to TRIzol homogenation and RNA was extracted. It was further subjected to DNAse treatment as per the protocol mentioned by TURBO DNA-free Kit (Invitrogen). Then for cDNA preparation,DNAse treated RNA samples were reverse transcribed by using Superscript II RNase H-reverse transcriptase (RT) through random hexamers as per the manufacturer’s instruction (Fermentas, Thermo Fisher Scientific,Waltham,MA,USA). The synthesised cDNA was stored at -20° C to be directly used for reverse transcriptase polymerase chain reaction (RT-PCR) .Specific oligonucleotides were used for the analysis of COX-2 (FP – 5’ AAAGGCCTCCATTGACCAGA -3’, RP 5’-GTGCTCGGCTTCCAGTATTG-3’), product size 373 bp and GAPDH (glyceraldehyde 3-phosphate dehydrogenase) (FP 5’-AGTGAGGAGCAGGTTGAGGA-3’ and (RP) 5’ -GAGGAGGGGAGATGATGTGA-3’ (Reverse primer), product size 244 bp were synthesized from Euro film, India. The PCR reactions were performed in 25ml reaction mixtures containing 2ml cDNA, 2.0 mM of dNTP, and 2.5ml of 10 x standard Taq reaction buffer, 1.0 unit of Taq DNA polymerase (New England BioLabs, Inc.) and 5μmol of each primer. The reactions were carried out in the Thermo Fischer MiniAmp thermal cycler. PCR steps for COX-2: initial denaturation at 94 °C for 5 min-1 cycle, followed by 35 cycles of 94 C for 45s, 55 C for 45 s, 72 C for 1 min and final extension at 72° C for 10 min. The steps for GAPDH: initial denaturation at 94°C for 3 min, followed by 35 cycles of 94°C for 30 s, 55°C for 30 s, 72°C for 1min and final extension at 72°C for 8 min. The amplified products were separated on 1.5 %agarose gel electrophoresis containing ethidium bromide. The intensity of COX-2 was determined using image quant Omega Flour TM (GEL Company, USA) using Omega Fluor Acquisition software. The expression of COX-2 was expressed as per cent band intensity relative to that of GAPDH. All RT-PCR experiments were performed in triplicate with the same results and the best result provided in the result section.

##### Statistical analysis

Results are expressed as Means ± SD with the p value<0.001 in volume, upper and lower circumference of both the groups. Statistical significance was analyzed with unpaired student’s T-test for comparison between the 2 groups day wise. The comparisons among the groups were performed through Graph pad (8.0.2) using one-way ANOVA (Tukey’s multiple comparison tests) to compare the mean values of each group with each other. *,p < 0.05, ***, p < 0.001, and ****, p < 0.0001 is significantly different from experimental control group

## 3. Results

### 3. A. Effect of CRR on tail morphology and kinetics

#### 1: Morphological changes

We carried out an experimental model of the post-acute -surgical based lymphedema. We observed that following the incision, the murine shown the acquired lymphatic swelling in both groups however the EC group (**fig 1**) showed more swelling. The hair follicles on the inflamed area were erected and the shiny texture was started to appear as the marked area started to swell. Inflammation area of the tail near the edema was hairless and this condition was continuous in both the upper and lower area of the edema area. These changes were less prominent in CRR group (**fig 2**)

**Figure 2:**
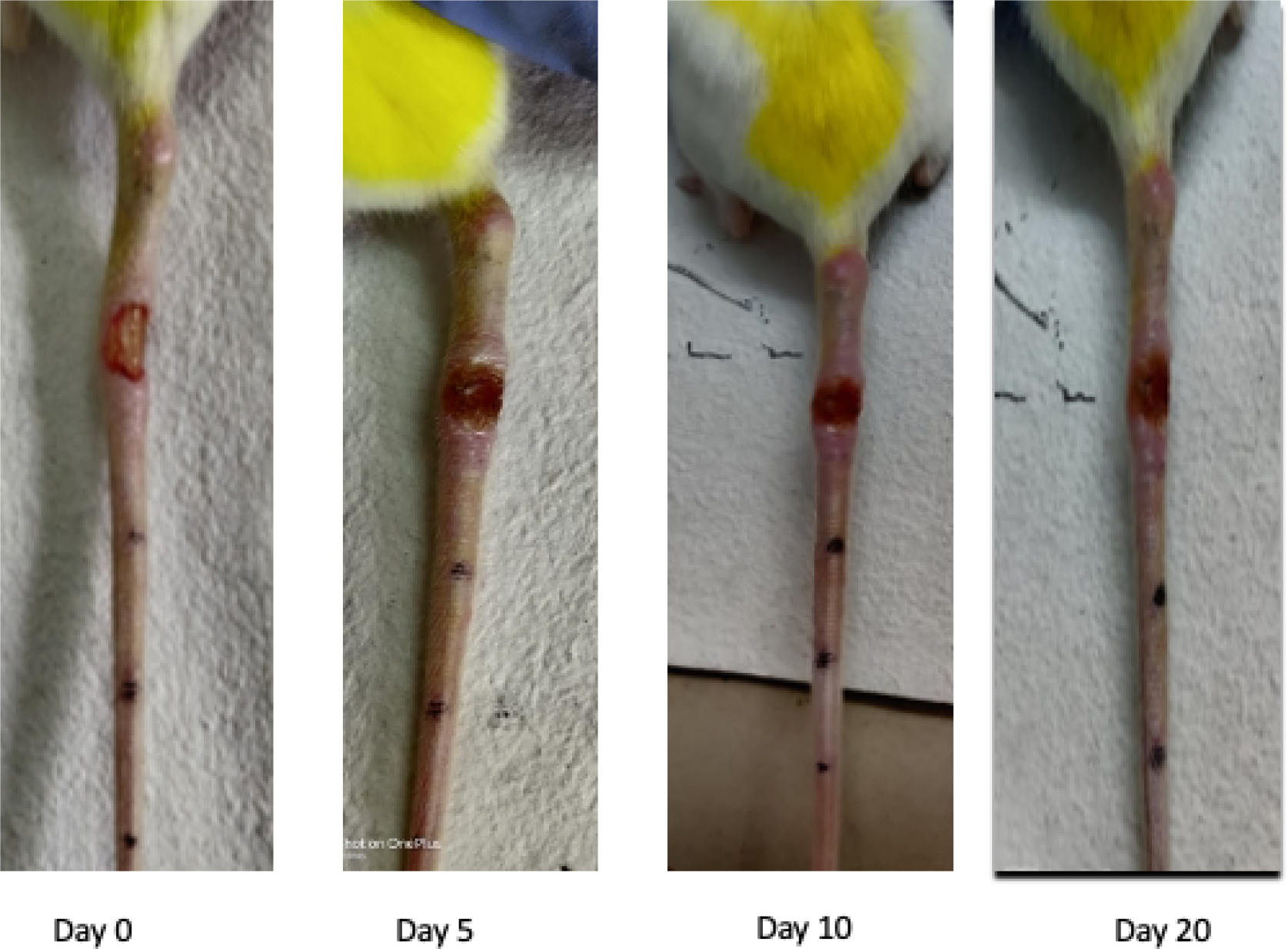
Morphological assessment of mice tail at day 0,5,10 and 20 in CRR group.

#### 2 Tail oedema post surgery changes

##### 1) On volume

The gradual increase of the tail’s total volume at post-surgical was observed using the formula mentioned in methodology. After the assessment of volume of the mice’s tail in both groups, the CRR group shown the significant decrease in volume compared to volume of the EC group. Significant difference (p<0.001) was observed from day 6 till end in comparison to EC group. and this difference was continued till the end point of the experiment. When observing the volume change, the most distinct increase was observed in day 11 where highest difference was observed. Though CRR group shown to attain the highest swelling area on the day 8 and day 9 but it was still low as compared to the EC which shown the highest swelling on the day 11^th^. Continuous decrease in volume was presented after day 9 for CRR and day 11 for EC. Furthermore, according to the assessment of the diameter the higher swelling was observed on the upper (Proximal) compared to lower (Distal) area in all the mice of the two groups. (**Figure 3 and 4)**

**Fig 3.**
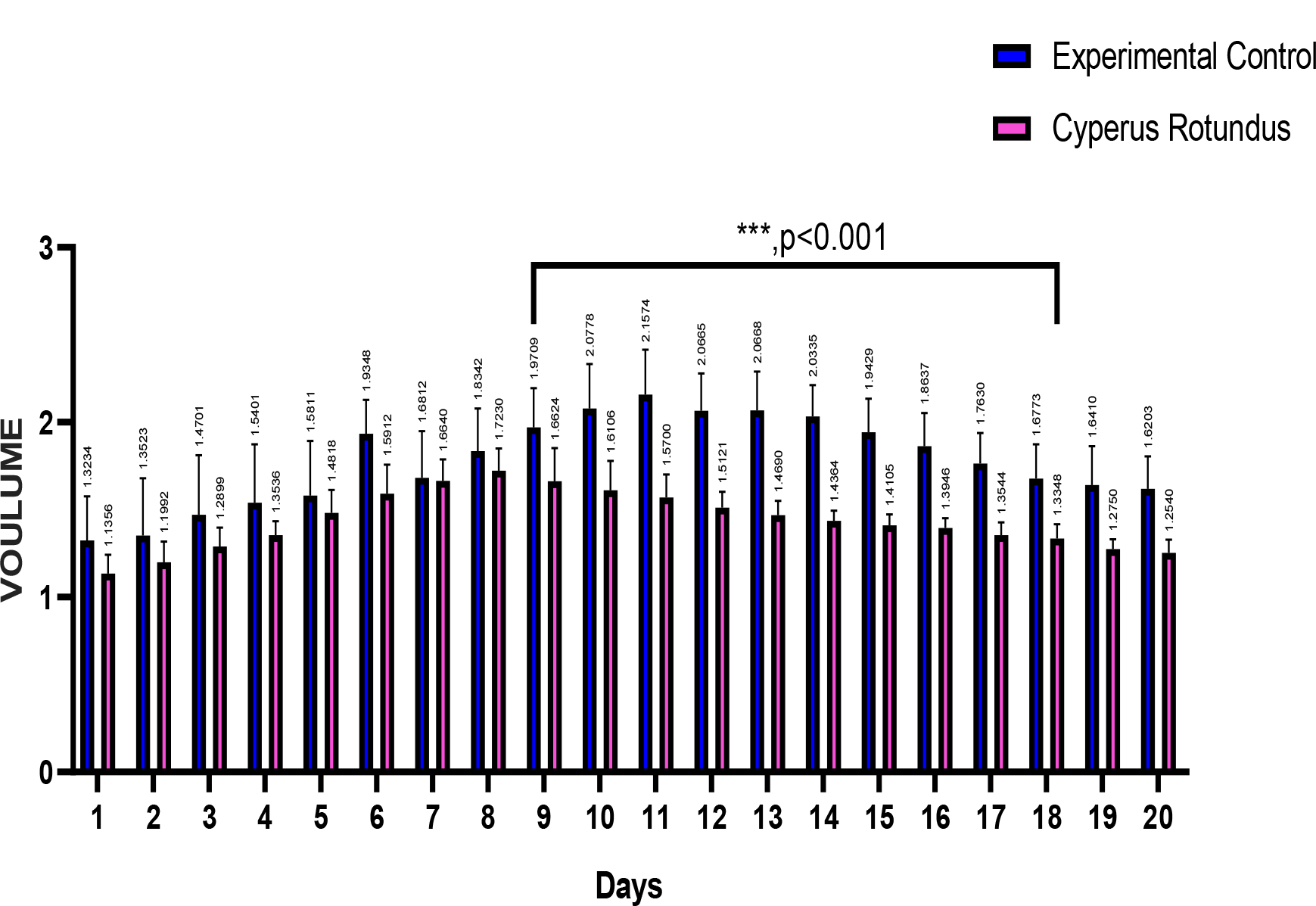
Graphical representation of the change in the volume of both groups. The Statistical analysis was done with Student’s t-test. The results were considered significant at _*p*_<0.001.

**Fig 4.**
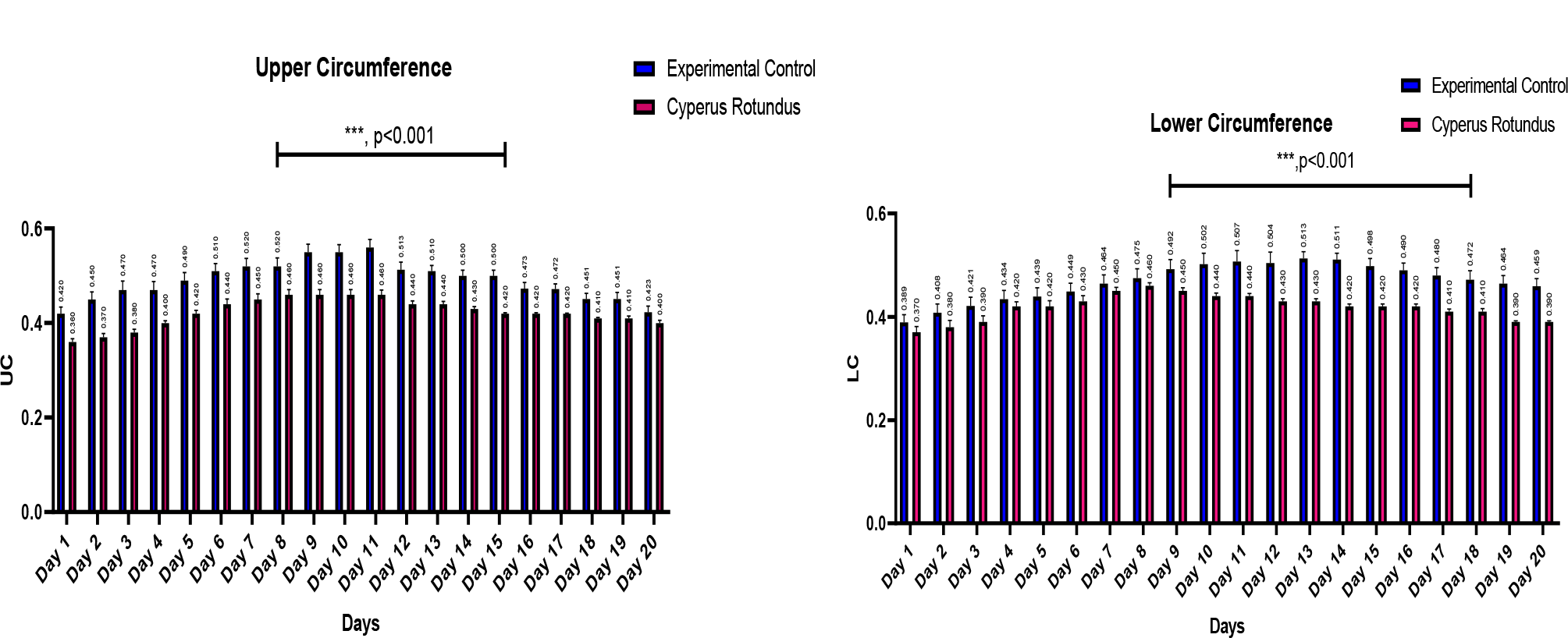
Graphical representation of the change in the circumference of the tail in both groups. Upper (a),lower(b) The Statistical analysis was done with Student’s t-test. The results were considered significant at _*p*_<0.001.

### 3. B. Effect of CRR on biochemical parameters

#### 1) On TNFα level

As the disruption to the lymphatic vasculature start the edema and inflammation. Hence to assess the inflammation in both the experimental control and *Cyperus rotundus* administered mice, inflammation was measured .The experimental group have shown the significant increment in the TNF-alpha, but in the case of CRR it was significantly lower in a mouse model of acquired lymphedema.(p<0.001) **(Fig 5**). Highest inflammation in the EC and CR were 170.73pg/ml and 148.221pg/ml respectively.

**Fig 5.**
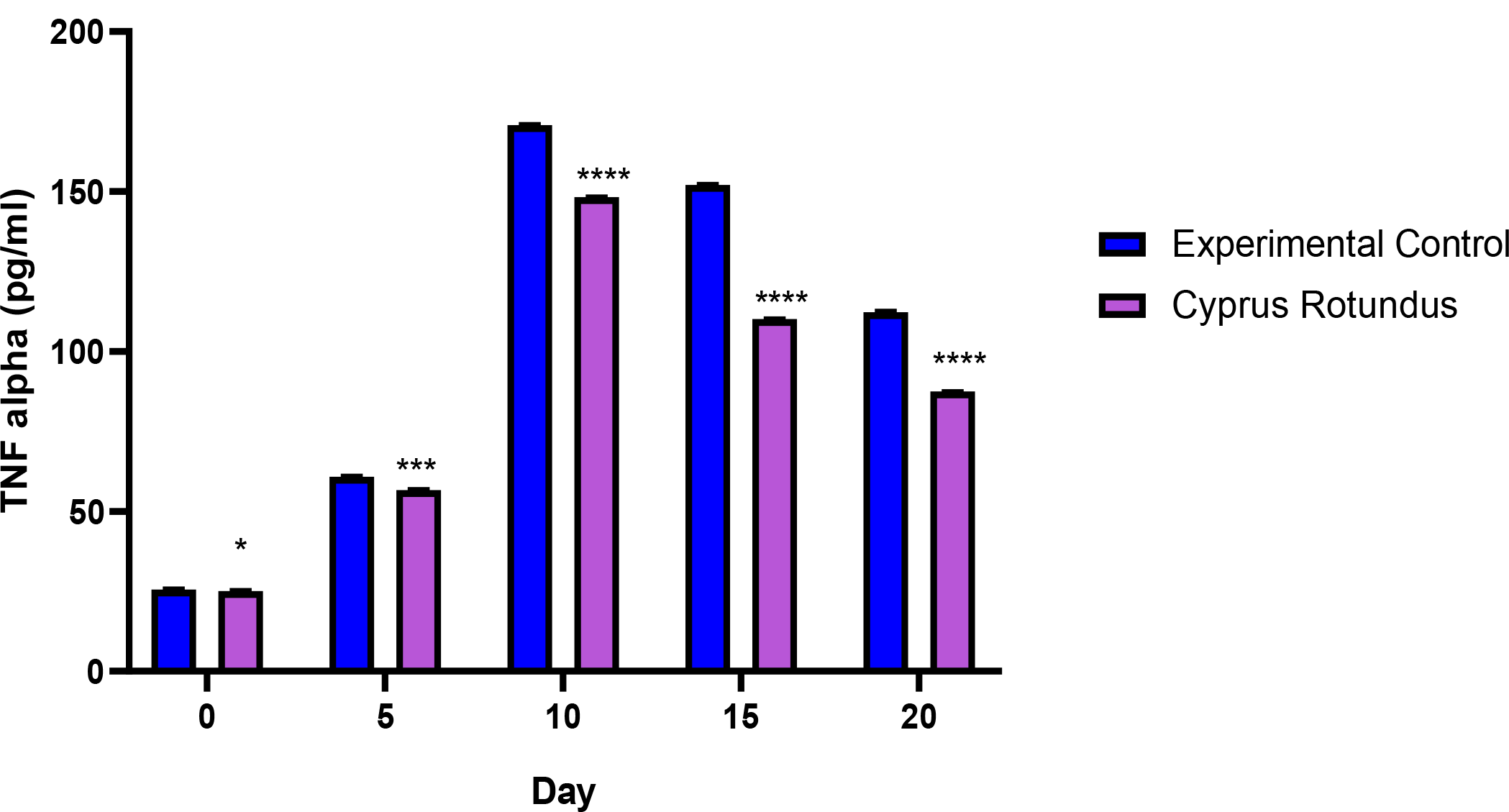
Level of TNFα in experimental control vs *Cyperus rotundus* at day 0, 5, 10, 15 and 20. Values represented as mean ± SD (n=5). The comparisons among the groups were performed using one-way ANOVA (Tukey’s multiple comparison tests) to compare the mean values of EC with CR. (^*^p < 0.05, ^***^p < 0.001, and ^****^p < 0.0001 is significantly different from the EC group in day wise.)

#### 2. On antioxidant enzyme status

The increased inflammation in the wounded area and its oxidative stress was determined by measuring antioxidant enzymes (SOD and CAT). It was observed that CRR group shown decreased SOD and CAT levels as compared to EC group .CRR treatment significantly elevated the SOD and CAT levels (p < 0.01) as equated to the EC group as evident from (**Figure 6 (a) (b)**).

**Fig 6.**
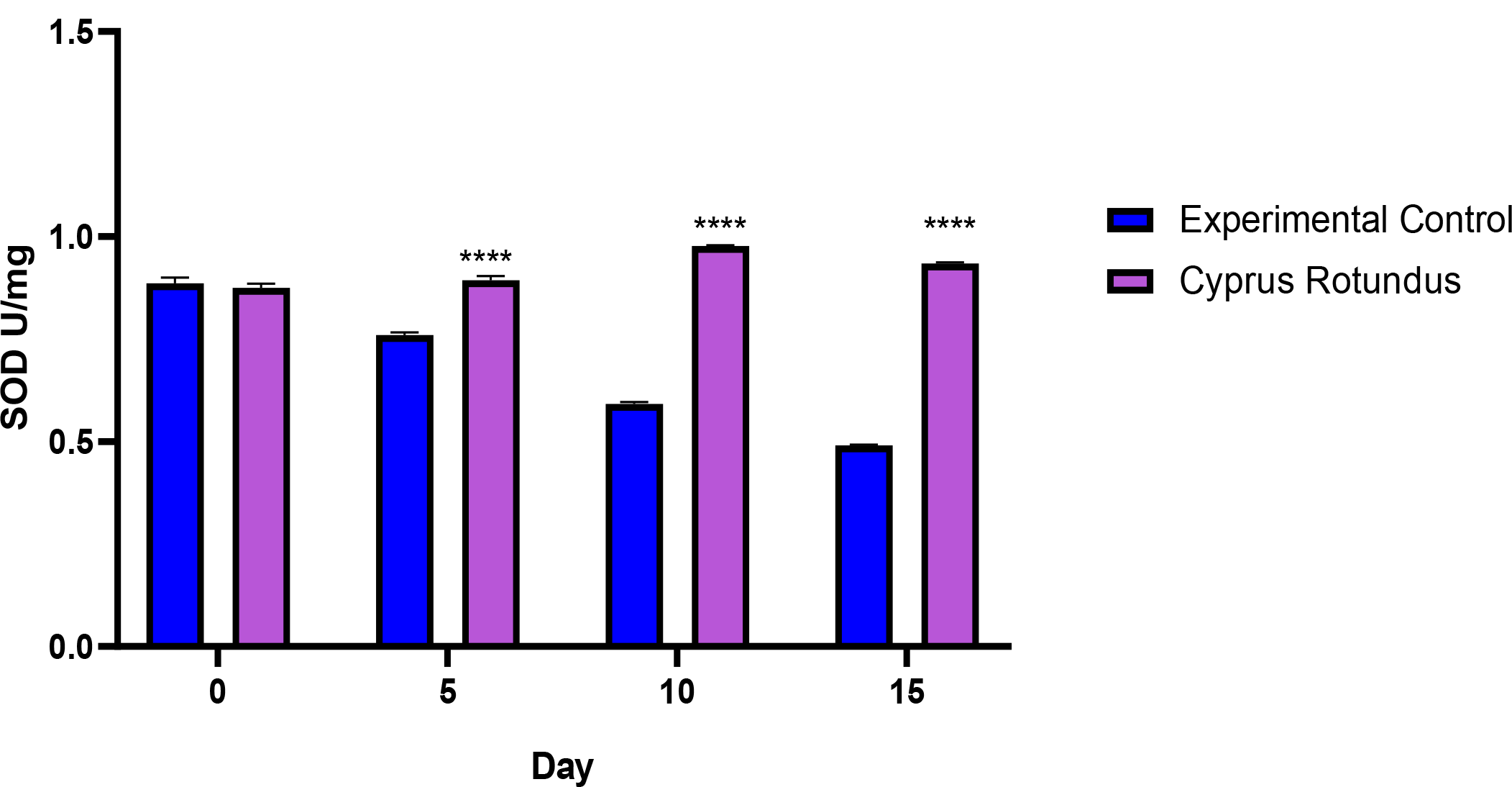
(a) SOD level in experimental control vs cyperus rotundus at day 0, 5, 10 and 15. Values represented as mean ± SD (n=5). The comparisons among the groups were performed using one-way ANOVA (Tukey’s multiple comparison tests) to compare the mean values of EC with CR. (^*^p < 0.05, ^***^p < 0.001, and ^****^p < 0.0001 is significantly different from the EC group day wise)

**Fig 6(b).**
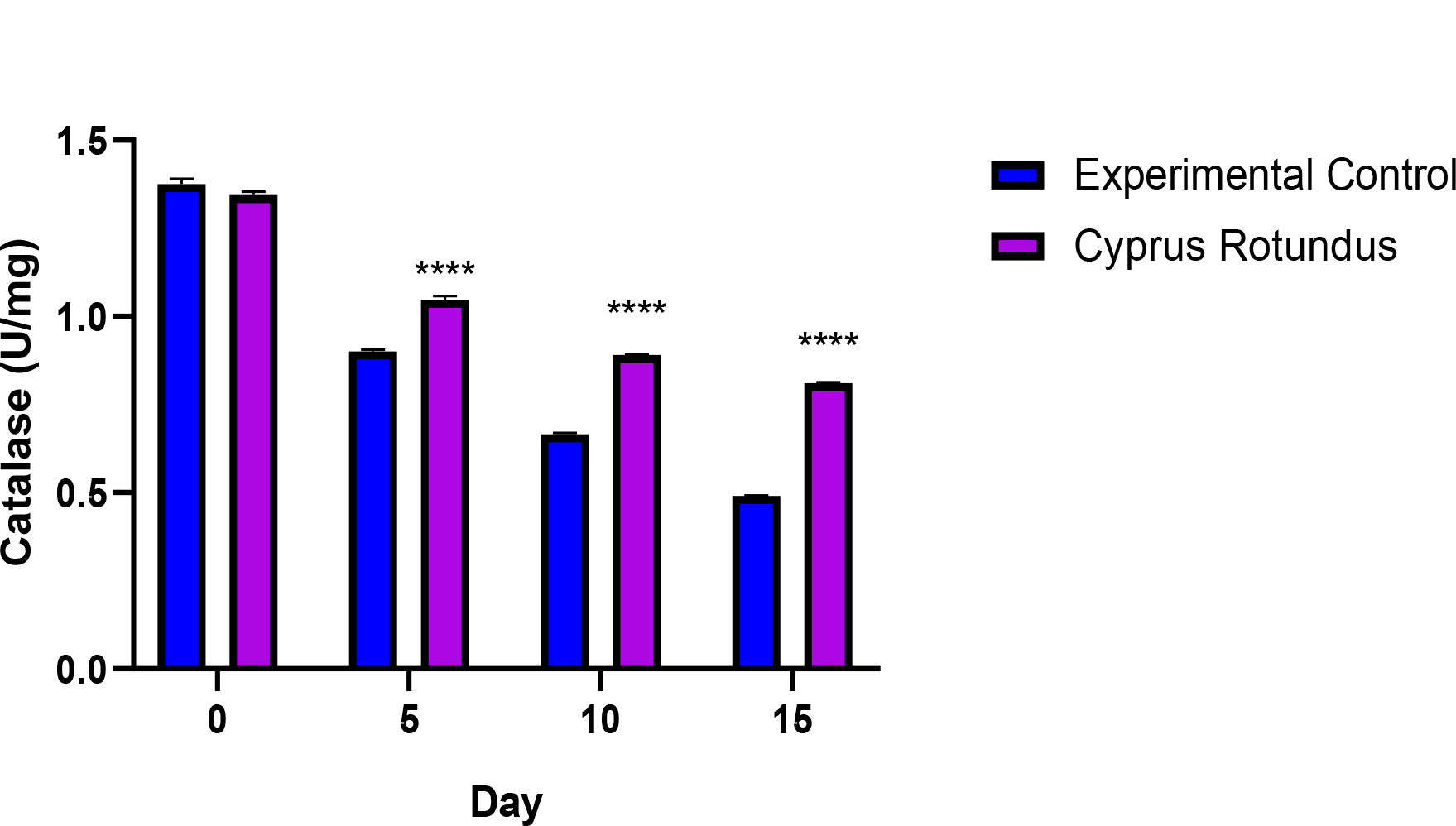
Catalase level in experimental control vs Cyperus rotundus at day 0, 5, 10 and 15. Values represented as mean ± SD (n=5). The comparisons among the groups were performed using one-way ANOVA (Tukey’s multiple comparison tests) to compare the mean values of EC with CR. (^*^p < 0.05, ^***^p < 0.001, and ^****^p < 0.0001 is significantly different from the EC group day wise)

**Fig 7.**
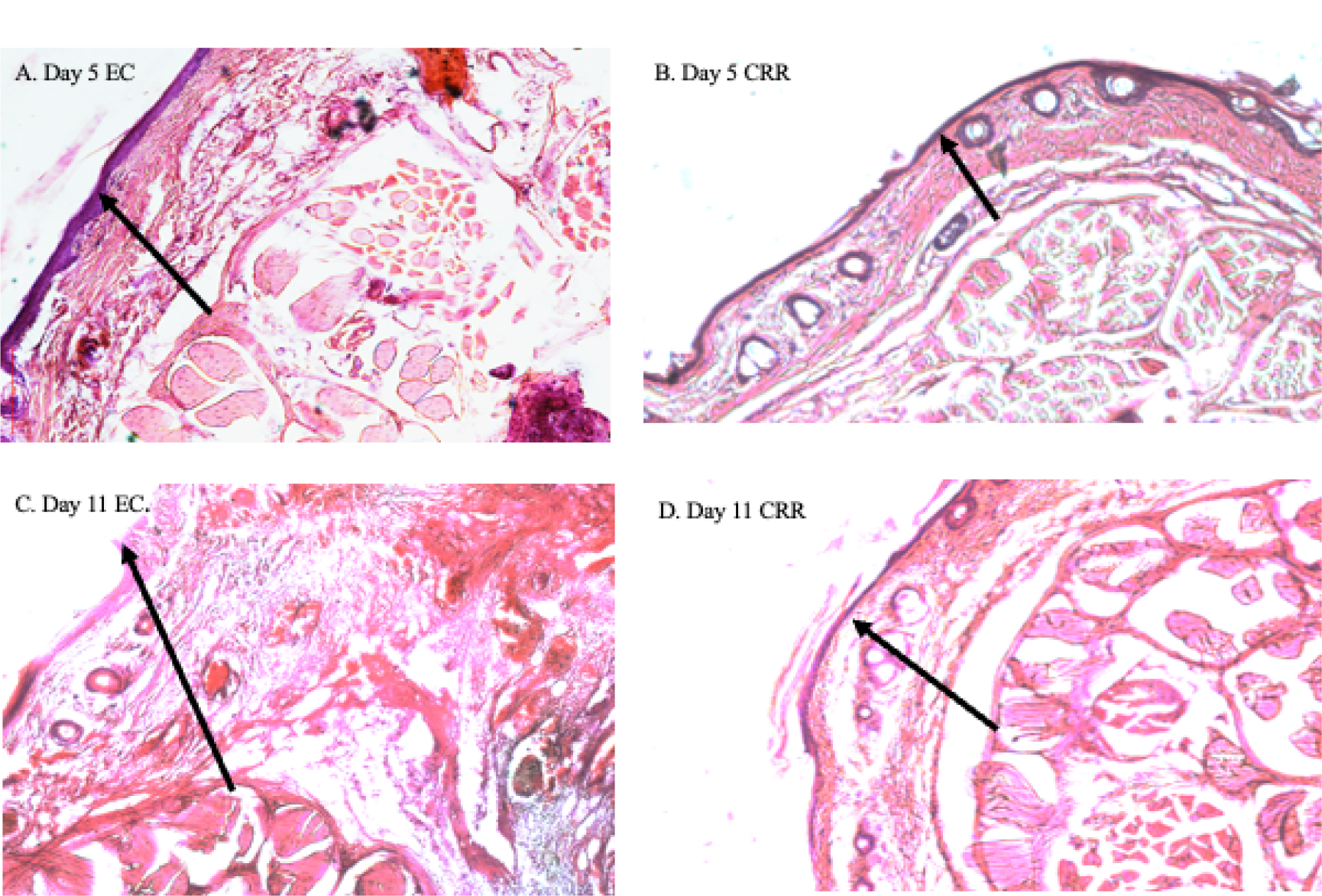
Histological observation (10X) in experimental control on day 5 (A) and day 11 (C) showed the thicker dermal, hypodermic perivascular and peri lymphatic layer. However in CRR treated group on day 5 (B) and day 11(D) there is reduced infiltration of lymphatic fluid and cells as compared to EC group.

**Fig 8.**
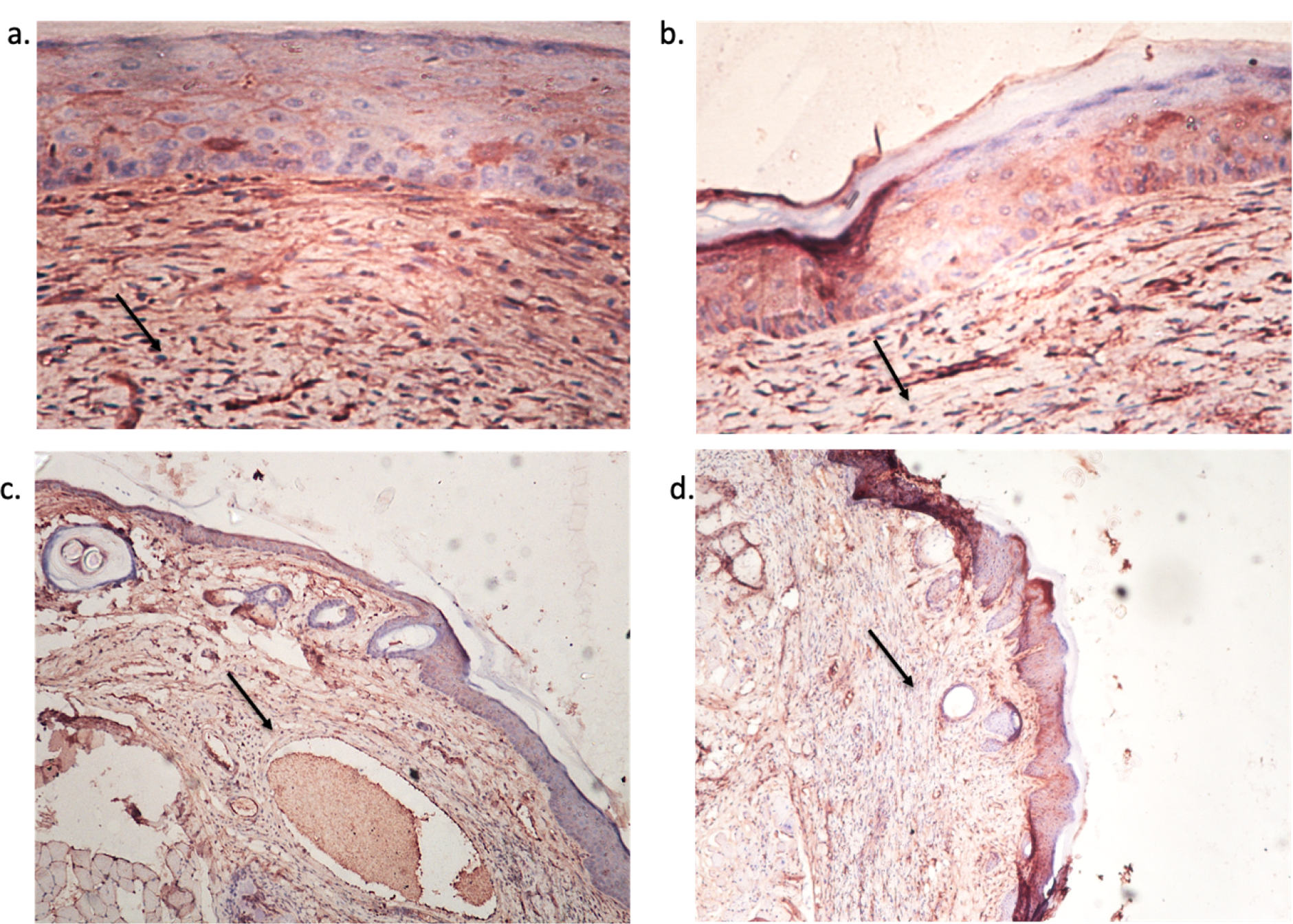
Immunostaining observation (10X) of CD 4 in the experimental control on day 5 (a) was observed high as compared to day 5 (b) of the CRR. In the day 11 (c) the pre-fasical lymphatics are dilated (arrow head) as compared to day 11 of CRR (d) and CD4 expression is higher in (c) as compared to (d).

**Fig 8 (a) (b).**
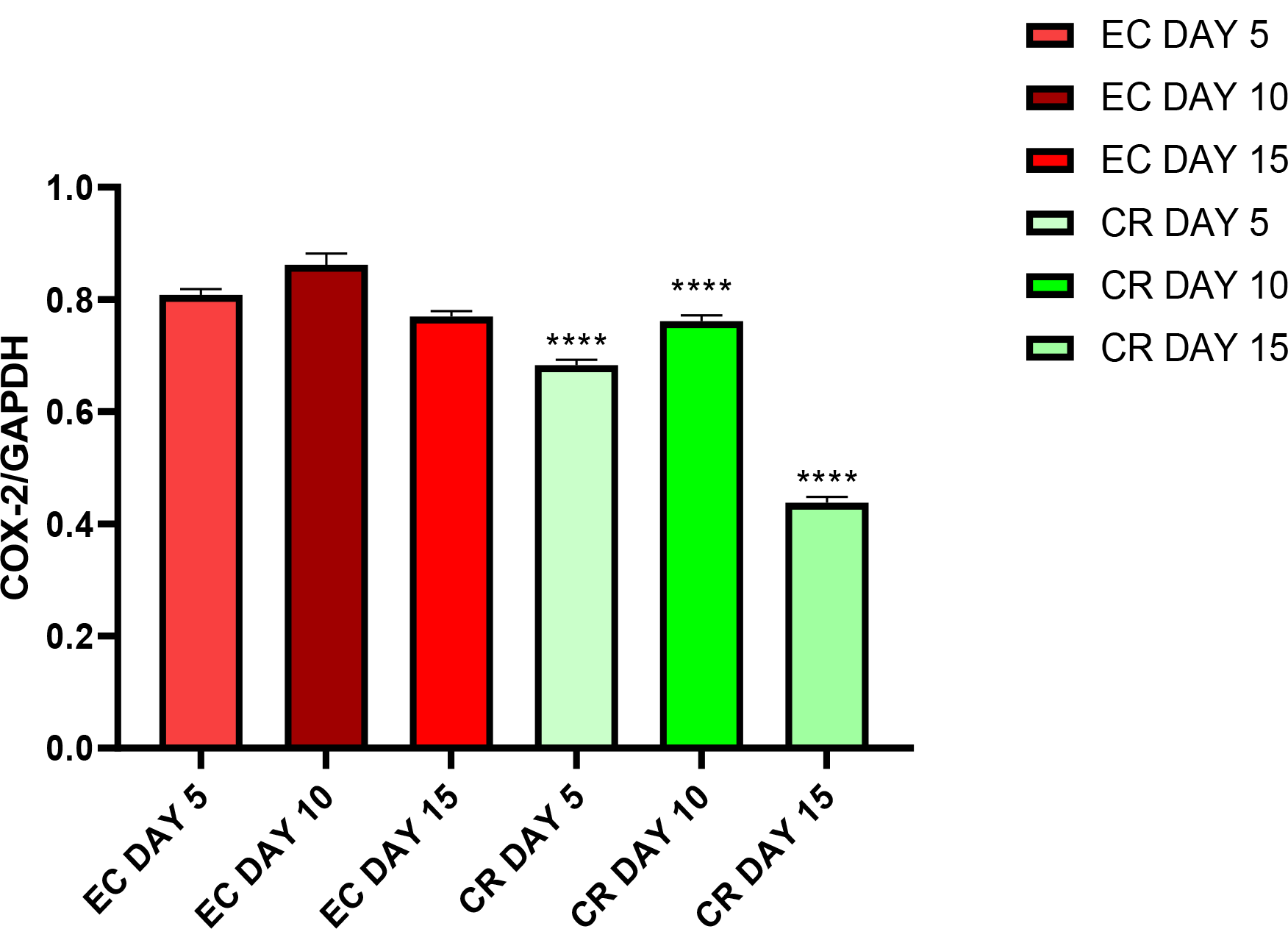
mRNA level of COX-2 in experimental control vs *Cyperus rotundus* at day 5, 10 and 15. Values represented as mean ± SD (n=3). The comparisons among the groups were performed using one-way ANOVA (Tukey’s multiple comparison tests) to compare the mean values of EC with CR. (^*^p < 0.05, ^***^p < 0.001, and ^****^p < 0.0001 is significantly different from the EC group day wise

### C. Effect of CRR on histological changes in mice tail

H&E staining of a cross section of the mouse tail shown to have different thickness in the EC tail in comparison to the CRR treated tail. While there are many infiltrated cells at the site of lymphedema in EC groups a lesser number of the inflammatory cells was observed in the CRR group (Fig.7(B)(D)).As the higher thickening in the dermal, perivascular and peri-lymphatic site can be seen EC group as compared to CRR administered group. Likewise, in EC group there was the marked inflammatory response, infiltration of leukocytes and higher collagen deposition in comparison to CRR group.

### D Effect of CRR on mRNA level of COX-2-

In the RT-PCR, the expression of the COX-2 was observed in both the EC and CR group, the findings indicate the up regulation of cox-2 in experimental group in EC Day 5, 10 and 15, in comparison to the *Cyperus rotundus* treated mice. In the day 15 we observed more down regulation of the expression in COX2 in both the EC and CR, but in comparison with day wise manner, CR group shown lower expression of cox-2 .In highest marked swelled area (DAY 11) of EC, the cox2 was highest but on the same day the CR group shown the lower expression of COX2 as well as in other parameters.

## DISCUSSION

Till date many research had been conducted on lymphedema but the molecular mechanism and targeted therapy remains debatable. The most common non-pharmacological treatment is decongestive physiotherapy which is used as supportive care for post management (16). Since the core question lies in the pathogenesis of lymphedema and its progression. It is an important domain to identify and utilise new therapeutic targets as currently there is no drug available in common person domain. The uncovering of the new compounds which can act on the various different pathways during the lymphedema is utmost important task for patients. As the secondary lymphedema induction in animals is utilised by researcher to understand the pathophysiology(17). How the therapeutic agent could be used to observe the effects of it on the site of the oedema is another important point for the study of SL .Towards this objective here for the first time we have validated the therapeutic response of Cyperus rotundus root extract (CRR) in mouse model of secondary lymphedema. In this work we evaluated the effect of ethanolic extract of CRR in terms of change in the tail oedema post-surgery, biochemical, morphological and molecular changes in comparison to experimental control.

As roots of CR claims to have good efficiency as anti-oxidant and anti-inflammatory agent, owing to presence of high flavonoids and polyphenols in polar and non-polar extract of CR(18). In silico studies indicated the effect of CRR as an inhibitor of 5-lypoxygenease and leukotriene A4 hydrolase (LTA4H) reported by Fenanir et al (19) .Thus it could be that the reduced swelling after post-surgery in CRR group could be due to the effect of cyperus the provides a insight attenuating swelling in mice tail can be supported by earlier reports from the in-silico studies indicating the role of CRR as Apart from this, another feature of lymphedema which is the deposition of adipose tissue which may triggered by subtle dysfunction or the injury in the lymphatic system. Cyperus rotundus root extract effect in the decrescent of swelling is strongly supporting its earlier described role as downregulating of pro-inflammatory cytokines genes. Molecule’s present in CR has been reported to inhibit the pro-inflammation via the inhibition of the NF-κB and STAT3 pathways and ROS. (20)(21).As in the condition of SL, some of the studies have shown the expression of the pro-inflammatory gene which are been upregulated in animal models and patients with lymphedema. Hence the expression of pro-inflammatory gene such as the TNF-alpha are seen to be at increased state(22).

Another factor is the increased ROS level which causes higher oxidative stress in the higher lymphedema fluid. These implications have been addressed in literatures on how the ROS are able to disturb the lymphatic contractions and how the robust anti-oxidant defence could play vital role in the secondary lymphedema. In the lymphedema, COX pathway sits in centre for the production of ROS in the surgically induced lymphedema site and due to increased ROS it also enhances lipid peroxidation .CRR have the potency to decrease the lipid peroxidation (23).

Therefore, due to high inflammation and increased ROS at the inflamed area brings the subcutaneous fat deposition and increased in fat thickness in the case of the mice tail skin which can be observed in histological analysis (24) (25) (26). Considering all these cascading effects arising after the induction of surgical based lymphedema, we propose that CRR can effectively downregulate the ROS level via inhibiting the COX -2 expression and also lowers the TNF alpha level. We utilised the mouse tail edema model to research this hypothesis .As shown in figures above. It was supported by the volume change in mice tail (Fig 3.) and change in circumferential area of both upper and lower area of surgical area also established this(Fig 4 (a)(b)).

Hence, from this study our results shows that CRR ethanolic extract shown its potency as anti-inflammatory and antioxidant property in the mice tail of lymphedema. Though more refined and single compounds based studies in future could aid in the management of secondary lymphedema and this could shed more light on the *Cyperus rotundus’s* derived compounds in the initiation of therapeutic use of it in future.

## Conclusions

Our work showed roots of *Cyperus rotundus* can be used to target secondary lymphedema by decreasing the level of inflammatory cytokines and reactive oxygen species. This was morphologically supported by change in the tail volume and circumference. We have a Swiss albino mice tail model for inducing secondary lymphedema, which can be used to stimulate a trait of human secondary lymphedema and provides the insights of functional and structural changes of lymphedema. Our work provides insight on the importance of COX-2 in development of secondary lymphedema and how CRR significantly reduced it after the post-surgery. However further studies are required for Cyperus rotundus to be a potential plant as an additional treatment option in future.

## Acknowledgement

NP and PM is thankful to ICMR and UGC respectively for providing the financial assistance.

